# An analysis and comparison of the statistical sensitivity of semantic similarity metrics

**DOI:** 10.1101/327833

**Authors:** Prashanti Manda, Todd Vision

## Abstract

Semantic similarity has been used for comparing genes, proteins, phenotypes, diseases, etc. for various biological applications. The rise of ontology-based data representation in biology has also led to the development of several semantic similarity metrics that use different statistics to estimate similarity.

Although semantic similarity has become a crucial computational tool in several applications, there has not been a formal evaluation of the statistical sensitivity of these metrics and their ability to recognize similarity between distantly related biological objects.

Here, we present a statistical sensitivity comparison of five semantic similarity metrics (Jaccard, Resnik, Lin, Jiang& Conrath, and Hybrid Relative Specificity Similarity) representing three different kinds of metrics (Edge based, Node based, and Hybrid) and explore key parameter choices that can impact sensitivity. Furthermore, we compare four methods of aggregating individual annotation similarities to estimate similarity between two biological objects - All Pairs, Best Pairs, Best Pairs Symmetric, and Groupwise.

To evaluate sensitivity in a controlled fashion, we explore two different models for simulating data with varying levels of similarity and compare to the noise distribution using resampling. Source data are derived from the Phenoscape Knowledgebase of evolutionary phenotypes.

Our results indicate that the choice of similarity metric along with different parameter choices can substantially affect sensitivity. Among the five metrics evaluated, we find that Resnik similarity shows the greatest sensitivity to weak semantic similarity. Among the ways to combine pairwise statistics, the Groupwise approach provides the greatest discrimination among values above the sensitivity threshold, while the Best Pairs statistic can be parametrically tuned to provide the highest sensitivity.

Our findings serve as a guideline for an appropriate choice and parameterization of semantic similarity metrics, and point to the need for improved reporting of the statistical significance of semantic similarity matches in cases where weak similarity is of interest

## 2 Introduction

Semantic similarity is a way of measuring partial or imperfect similarity between objects based on the similarity of their ontology annotations Pesquita et al. [2009]. In biology, semantic similarity has been used to compare proteins, phenotypes, diseases, and other biological objects Jain and Bader [2010]; Robinson et al. [2014]; Washington et al. [2009]; Manda et al. [2015]. For example, semantic similarity is the cornerstone for hypothesizing candidate genes for evolutionary transitions in the Phenoscape project Edmunds et al. [2015]; Manda et al. [2015] and for connecting model organism phenotypes to human disease phenotypes for rare disease diagnosis within the Monarch Initiative Mungall et al. [2015, 2016]. In some applications, such as in Phenoscape, the biological objects being compared are distantly related requiring semantic similarity approaches that can effectively identify low degrees of partial relatedness Manda et al. [2015].

Although semantic similarity comparisons have become the linchpin of several applications, and, are often used to compare distantly related biological objects, it is unclear how sensitive these metrics are to the degree of similarity. It is also unknown if sensitivity to imperfect matches differs among different similarity metrics. In addition to the choice of a similarity metric, there are several other parameters that influence how similarity between biological objects is computed and may influence sensitivity. One is how individual similarities between two objects’ annotation sets are summarized to determine the similarity between the objects. Another is the difference between symmetric metrics versus asymmetric metrics.

Here, we conduct a statistical sensitivity analysis of a set of semantic similarity metrics combined with different choices of parameters to assess robustness at identifying different levels of imperfect similarity. This sensitivity analysis can serve as a guide for evaluating different semantic similarity metrics, and relevant parameters to enable appropriate choices for any particular application.

First, we introduce an application within the Phenoscape project that requires the comparison of distant biological objects (Manda et al. [2015, 2016]) and use it for evaluating the sensitivity of semantic similarity metrics and associated parameters.

Phenotypic diversity among species is typically described in phylogenetic studies using characters consisting of two or more states that vary in some attribute of that character, such as size or shape Dahdul et al. [2010]; Mabee et al. [2012]. These evolutionary character states are associated with taxa at different levels in a phylogenetic tree. Species closely related to each other will generally have more character states in common Dahdul et al. [2010]. Model organism communities have studied phenotypes, but in this case resulting from genetic perturbations of specific genes Sprague et al. [2007]; Smith et al. [2005]; Karpinka et al. [2015]. Connecting evolutionary variation among taxa on a phylogenetic tree to knowledge from developmental genetics using semantic similarity can expand our understanding of both evolutionary variation and genetics, and this is a major goal of the Phenoscape project Manda et al. [2015]; Dahdul et al. [2010]; Edmunds et al. [2015]; Mabee et al. [2012]. The Phenoscape Knowledgebase (KB) currently contains 415,819 phenotype annotations that correspond to 21,570 evolutionary character states in 5,211 vertebrate taxa, 4,161 of which are terminal. These are integrated with phenotypes for 3,526 human, 7,758 mouse, 5,883 zebrafish, and 12 *Xenopus* genes from the Human Phenotype Ontology Köhler et al. [2013], Mouse Genome Informatics, Eppig et al. [2015], Zebrafish Information Network Bradford et al. [2011], and Xenbase Karpinka et al. [2015], respectively, and aims to support semantic similarity matching among phenotypes within and between these different sources Manda et al. [2015].

However, there are several challenges in comparing evolutionary phenotypes to gene phenotypes using semantic similarity. First, evolutionary and model organism gene phenotypes are studied by different research communities who describe them using different ontologies and different annotation formats. For example, evolutionary phenotypes are often annotated in the Entity Quality (EQ) annotation format Mungall et al. [2010]. The Entity (*e.g.* ‘anal fin’) is drawn from one ontology, such as the animal anatomy ontology UBERON Haendel et al. [2014], and the Quality (*e.g.* ‘elongated’), that describes how the Entity is affected, is drawn from a trait ontology, such as PATO Gkoutos et al. [2005]; Mungall et al. [2010]. In contrast, model organism gene phenotypes such as ones from mouse are described using the Mammalian Phenotype ontology Smith et al. [2005]. Second, the species and their anatomical structures being described in evolutionary and model organism phenotypes can be vastly different. Even when changes in the same genetic pathways affect the same anatomical structures, the phenotypes that have changed over evolution will generally be different from those induced in the laboratory. Given these various considerations, exact matches between phenotypes from these different data sources will be vanishingly rare. It is essential that semantic similarity measures have the ability to detect very weak matches, and that one can recognize when the best match available is sufficiently weak that it cannot be distinguished from noise.

There are several important parameter choices in the computation of semantic similarity that can affect sensitivity. First, we address choice of the metric itself. Several semantic similarity measures have been developed and applied to biological data Pesquita et al. [2009]. Some of these metrics primarily use ontology structure while others also use statistics such as Information Content (IC) computed from annotation data Pesquita et al. [2009].

The set of ontology annotations used to describe an object is said to be the object’s profile. After computing similarity between two sets of individual annotations, there are several approaches to aggregate the individual similarities to estimate similarity between the two objects. For example, the All Pairs profile similarity approach uses the distribution of all pairwise similarities between annotations in the two profiles of objects being compared Pesquita et al. [2009]. In contrast, the Best Pairs approach considers only the distribution of best matches for annotations in a profile Pesquita et al. [2009]. In both approaches, different representations of the above similarity distribution such as the mean, median, or a different quantile can be used to quantify similarity between the objects being compared. In contrast to All Pairs and Best Pairs, Groupwise aggregation determines similarity between two objects by computing set-based operations over the two profiles Pesquita et al. [2009]. Another issue to consider for semantic similarity computation is that some semantic similarity metrics for profiles, such as Best Pairs, are not commutative, which is to say that the similarity between object *A* to object *B* may differ from that of *B* to *A*.

Since it is not clear how these choices affect the sensitivity for detecting weak semantic similarities, we sought to address this within the context of our application of interest - comparison of biological phenotypes. We designed a series of experiments to assess the sensitivity of a set of semantic similarity metrics, based on how well the measure of similarity between increasingly imperfect profiles could be distinguished from noise. We introduce two models for increasing dissimilarity between initially identical phenotypes, and simulate annotations under these models using sourse data from the Phenoscape KB.

### 2.1 Semantic similarity metrics

A variety of semantic similarity metrics have been developed and applied to compare biological entities Pesquita et al. [2009]. These metrics can be broadly classified into edge-based, node-based, and hybrid measures Pesquita et al. [2009]. Edge-based metrics primarily use distance between terms in the ontology as a measure of similarity. Node-based measures use Information Content of the terms being compared and/or their least common subsumer (LCS). Hybrid measures incorporate both edge distance and Information Content to estimate similarity between ontology terms. Jaccard (edge-based) and Resnik (node-based) similarity are two of the most widely used similarity metrics for biological applications Washington et al. [2009]. We selected Jaccard from the edge based category and Resnik, Lin, Jiang, and Conrath from the node based category Pesquita et al. [2009]. From the set of hybrid measures, Hybrid Relative Specificity Similarity (HRSS) Wu et al. [2013] was selected because this metric was shown to outperform other metrics in tests such as distinguishing true proteinprotein interactions from randomized ones and obtaining the highest functional similarity among orthologous gene pairs Wu et al. [2013].

## 3 Methods

### 3.1 Simulating decay of semantic similarity

A database of 659 simulated evolutionary profiles with the same size distribution as the 659 true profiles in the Phenoscape KB was created by selecting annotations with uniform probability and without replacement from the pool of evolutionary annotations. Five query profiles each of sizes 10, 20, and 40 were randomly selected from the simulated database. Three different profile sizes were examined.

Next, “decayed” profiles were created for each simulated profile using one of the two models described below, in order to compare the query profile to progressively dissimilar profiles. Initially, the query profile is a perfect matches to itself, but similarity eventually decays until it is no longer distinguishable from noise. To characterize the noise distributions, we also generated 5,000 profiles of the same size as the query by drawing annotations randomly from among the 57.51 available. These profiles would not be expected to have more than nominal similarity with any of the simulated subject profiles.

These two decay models reflect two different ways in which we might simulate semantic similarity progressively decreasing between two profiles.

#### 3.1.1 Decay by Random Replacement

In the Decay by Random Replacement (*R*_*R*_) approach (Figure 1), annotations in the query profile are replaced, one per iteration, by an annotation selected randomly, with replacement, from the pool of 57.51 annotations. The process terminates when all annotations in the profile have been replaced. Thus, there is a 1-step decayed profile in which one original annotation has been replaced, a 2-step decayed profile in which two have been replaced, and so on.

**Figure 1:**
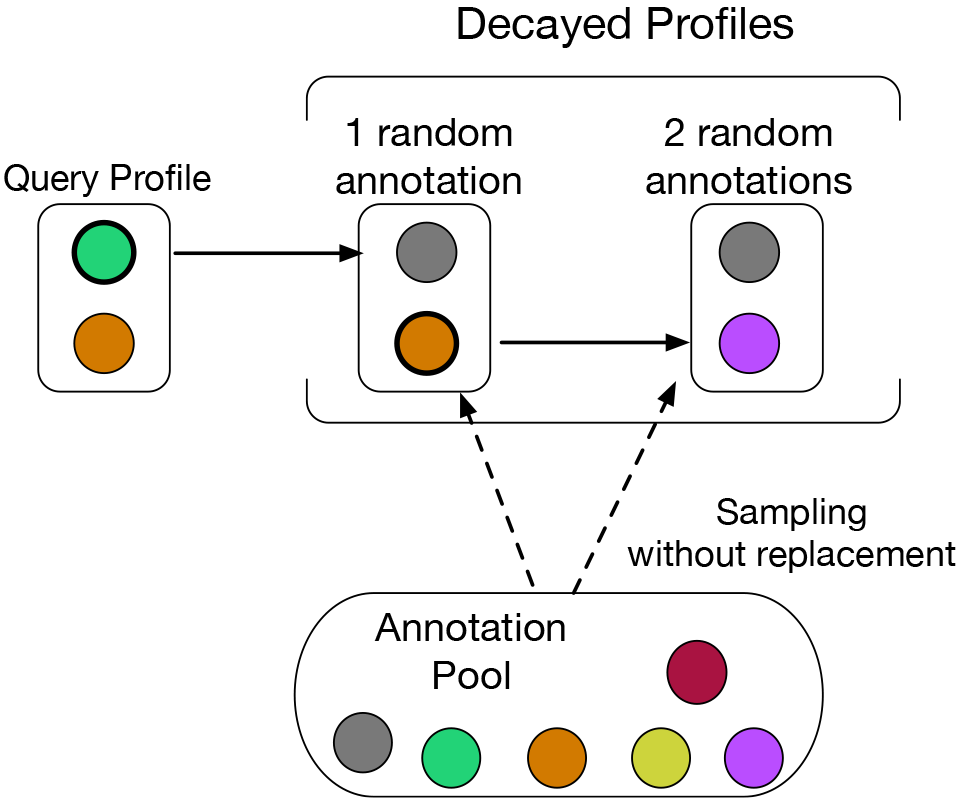
Profile decay via random replacement. Profiles are decayed iteratively with one annotation per iteration being replaced with a random selection from the pool of all possible annotations. The decay process ends when every original annotation in the profile has been replaced.

#### 3.1.2 Decay by Ancestral Replacement

In the Decay by Ancestral Replacement (*R*_*A*_ approach (Figure 2), annotations in the query profile are replaced, one per iteration, by progressively more semantically distant sibling, cousin, or parent terms. If an annotation has no siblings, it is replaced by an immediate parent. If an annotation has multiple immediate parents, a parent is selected randomly from the set of parents for replacement. Unlike the *R*_*R*_ approach which only goes through *N* replacements for a profile of size *N*, the *R*_*A*_ approach continues the decay process after each annotation in the profile has been replaced once. Subsequent iterations further decay the modified profile from the previous iteration by replacing each annotation with a more distantly related term. The process of decaying the query profile can be terminated when all annotations in the profile converge at the ontology root or when there is no further decay in similarity.

**Figure 2:**
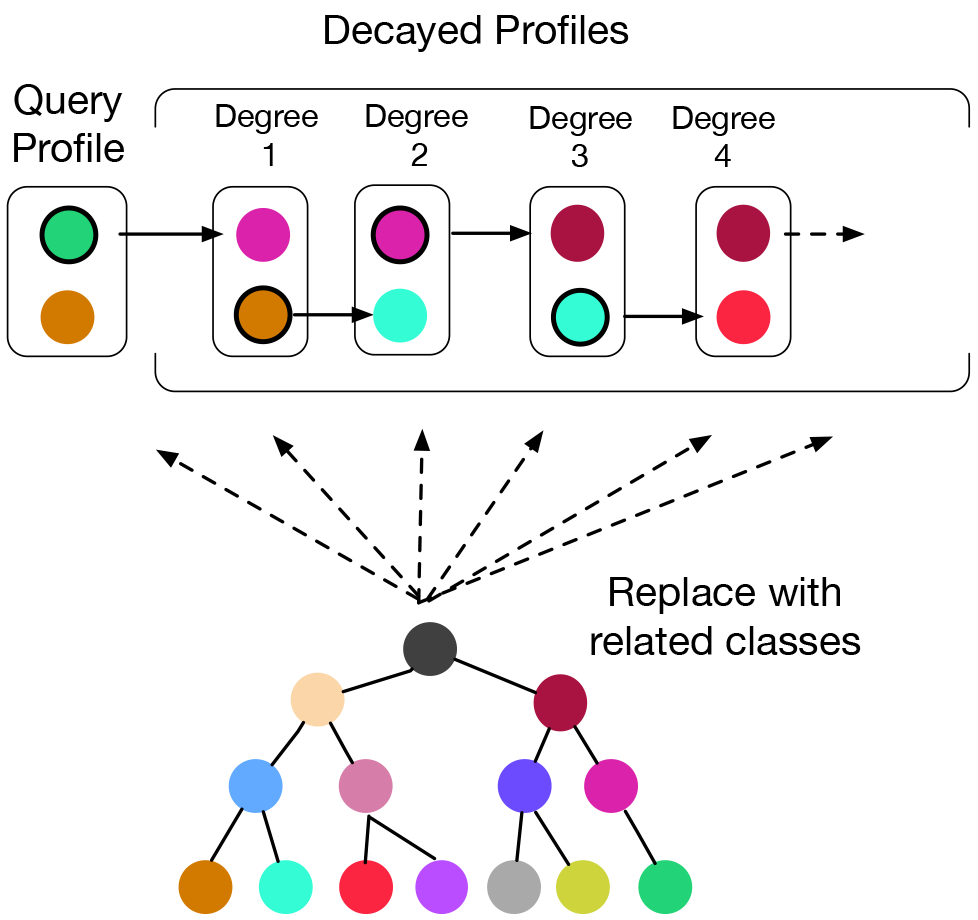
Profile decay via ancestral replacement. Profiles are decayed iteratively with one annotation per iteration being replaced with a related ontology class (sibling, cousin, parent). The decay process ends the process converges at the root or when there is no further decay in similarity.

#### 3.2 Semantic similarity metrics

##### 3.2.1 Jaccard similarity

The Jaccard similarity (*s*_*J*_) of two classes (*A*, *B*) in an ontology is defined as the ratio of the number of classes in the intersection of their subsumers over the number of classes in their union of their subsumers Mistry and Pavlidis [2008].

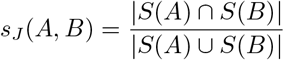
where *S*(*A*) is the set of classes that subsume *A*.

##### 3.2.2 Resnik similarity

The Information Content of ontology class *A*, denoted *I*(*A*) is defined as the negative logarithm of the proportion of profiles annotated to that class *f*(*A*) out of *T* profiles in total.

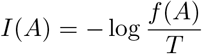

Since the minimum value of *I*(*A*) is zero at the root of the ontology, while the maximum value is −log(1/*T*), we can compute a Normalized Information Content (*I*_*n*_) with range [0, 1]

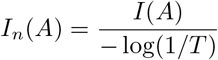

The Resnik similarity (*s*_*R*_) of two ontology classes is defined as the Normalized Information Content of the least common subsumer (LCS) of the two classes.

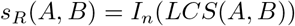

##### 3.2.3 Jiang and Conrath

Jiang and Conrath similarity (*s*_*C*_) takes into account the IC of two ontology classes as well as the IC of their Least Common Subsumer (LCS) Gan et al. [2013].

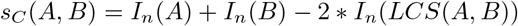

##### 3.2.4 Lin

Lin similarity (*s*_*L*_) also takes into account the IC of the two ontology classes and the IC of their least common subsumer (LCS), but in a different way Gan et al. [2013].

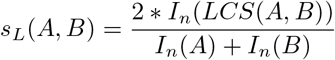

##### 3.2.5 Hybrid Relative Specificity Similarity

Hybrid Relative Specificity Similarity (HRSS, *s*_*H*_(*A*, *B*)) combines edge based and IC based measures. HRSS takes into account the specificity of the classes being compared along with their generality by using the LCS and the Most Informative Leaves (MIL) of the classes Wu et al. [2013].

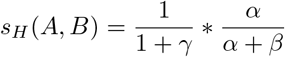

where

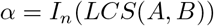

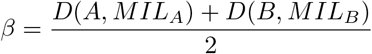

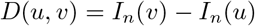

where *MIL*_*i*_ is the MIL of class *i*, *u* and *v* are ontology terms, and *u* is an ancestor of *v*.

### 3.3 Profile similarity

An annotation profile may consist of several annotations to a single object, such as a taxon or gene. In order to provide a single measure of the similarity of two objects when there are multiple pairwise similarity measures available between individual annotations, several methods are commonly used.

#### 3.3.1 All pairs

The All Pairs score (*a*_*z*_) between two entities *X* and *Y* (*a*_*z*_(*X*,*Y*)) is calculated by computing all pairwise annotation similarities between the annotation sets of *X* and *Y*. These |*X*|*|*Y*| pairwise annotation similarities can be summarized by taking the median or another quantile (or summary measure).

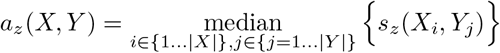

The index *z* can be used to specify the semantic similarity metric used in the computation.

#### 3.3.2 Best Pairs

To compute the Best Pairs score (*b*_*z*_) between *X* and *Y*, for each annotation in *X*, the best scoring match in *Y* is determined, and the median of the |*X*| resultant values is taken (*b*_*z*_(*X*,*Y*)).

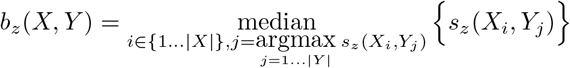

Unlike All Pairs, Best Pairs is not a commutative measure. To address this, a symmetric version (*p*_*z*_(*X*, *Y*)) of Best Pairs can be used.

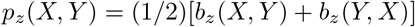

#### 3.3.3 Groupwise

Groupwise approaches (*g*_*z*_) compare profiles directly based on set operations or graph overlap. The Groupwise Jaccard similarity of profiles *X* and *Y*, *g*_*J*_ (*X*, *Y*), is defined as the ratio of the number of classes in the intersection to the number of classes in the union of the two profles.

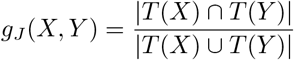

where *T*(*X*) is the set of classes in profile *X* plus all their subsumers.

Similarly, the Groupwise Resnik similarity of profiles *X* and *Y*, *g*_*R*_(*X*,*Y*), is defined as the ratio of the normalized Information Content summed over all nodes in the intersection of *X*, *Y* to the Information Content summed over all nodes in the union.

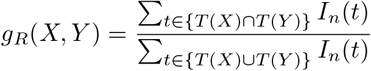

where *T*(*X*) is defined as above.

## 4 Results

The Phenoscape Knowledgebase contains a dataset of 661 taxa with 57,051 evolutionary phenotypes, which are phenotypes that have been inferred to vary among the taxon’s immediate descendents Manda et al. [2015]. A simulated dataset of subject profiles having the same size distribution of annotations per taxon was created by permutation of annotations.

### 4.1 Effect of query profile size on similarity decay

First, we wanted to determine how the pattern of similarity decline varies between different query profile sizes. We randomly selected five simulated query profiles each of size 10, 20, and 40 and created decayed profiles using the Decay by Random Replacement *(R_*R*_)* approach. The query profiles along with their decayed profiles were compared to the simulated database and the similarity score of the best match was plotted. Similarity was computed using five semantic similarity metrics (Jaccard, Resnik, Lin, Jiang, and HRSS) along four profile similarity settings (All Pairs, Best Pairs, Best Pairs Symmetric, and Groupwise (Jaccard and Resnik only)).

The results (Figure 3) indicate that the pattern of similarity decay is very similar across the three query profile sizes. This trend was observed consistently across different similarity metrics and profile similarity choices. For the Best Pairs methods, a sharp decline in similarity is observed at the 50% decay mark for all profile sizes. Groupwise metrics show a pattern of gradual decline across the three profile sizes. Given that pattern of decay was very similar across query profile sizes for all metrics, we only used query profiles of size 10 for the rest of the experiments in this study.

**Figure 3:**
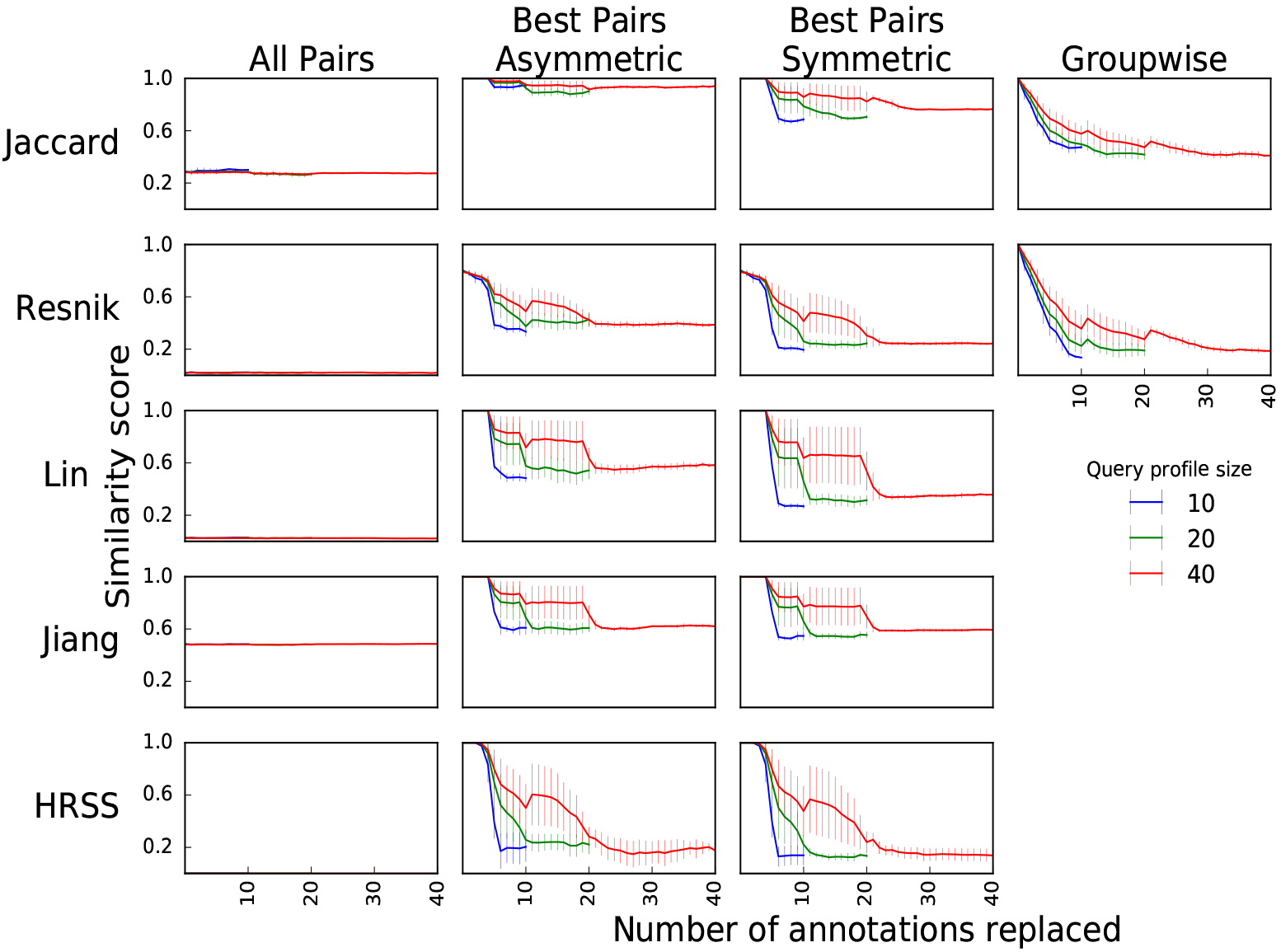
Decline in similarity for five simulated query profiles and associated decayed profiles each of size (blue), 20 (green), and 40 (red). Solid lines represent the mean best match similarity of the five query profiles to the database after each annotation replacement. Error bars show two standard errors of mean similarity across the five profiles.

### 4.2 Effect of profile similarity method on sensitivity of similarity metrics

Next, we conducted two comparisons - the four profile similarity approaches against each other, and the five semantic similarity metrics against each other. For ease of interpretation, we take the upper 99.9% of the similarity distribution for random profile matches (noise) as an arbitrary threshold for comparing the sensitivity of the different series. We use discrimination of similarity from the noise threshold as an indicator of the sensitivity of metrics. We observed discrimination from noise on two factors - the magnitude of discrimination (the higher, the better), and the point in the decay process at which similarity was no longer distinguishable from noise (the higher, the better).

The All Pairs profile similarity method (Figure 4, Col 1) fails to distinguish match similarity from noise across all five metrics. Both the Best Pairs variants (Figure 4, Cols 2, 3) demonstrate an initial discrimination from the noise threshold followed by a sharp decline around the 50% decay mark after which similarity falls below the noise threshold. The Symmetric measure (Figure 4, Col 3) shows substantially greater discrimination from noise as compared to the Asymmetric measure. Thus, both Best Pairs variants show greater sensitivity as compared to All Pairs, the Best Pairs Symmetric performing the best among the two Best Pairs methods.

**Figure 4:**
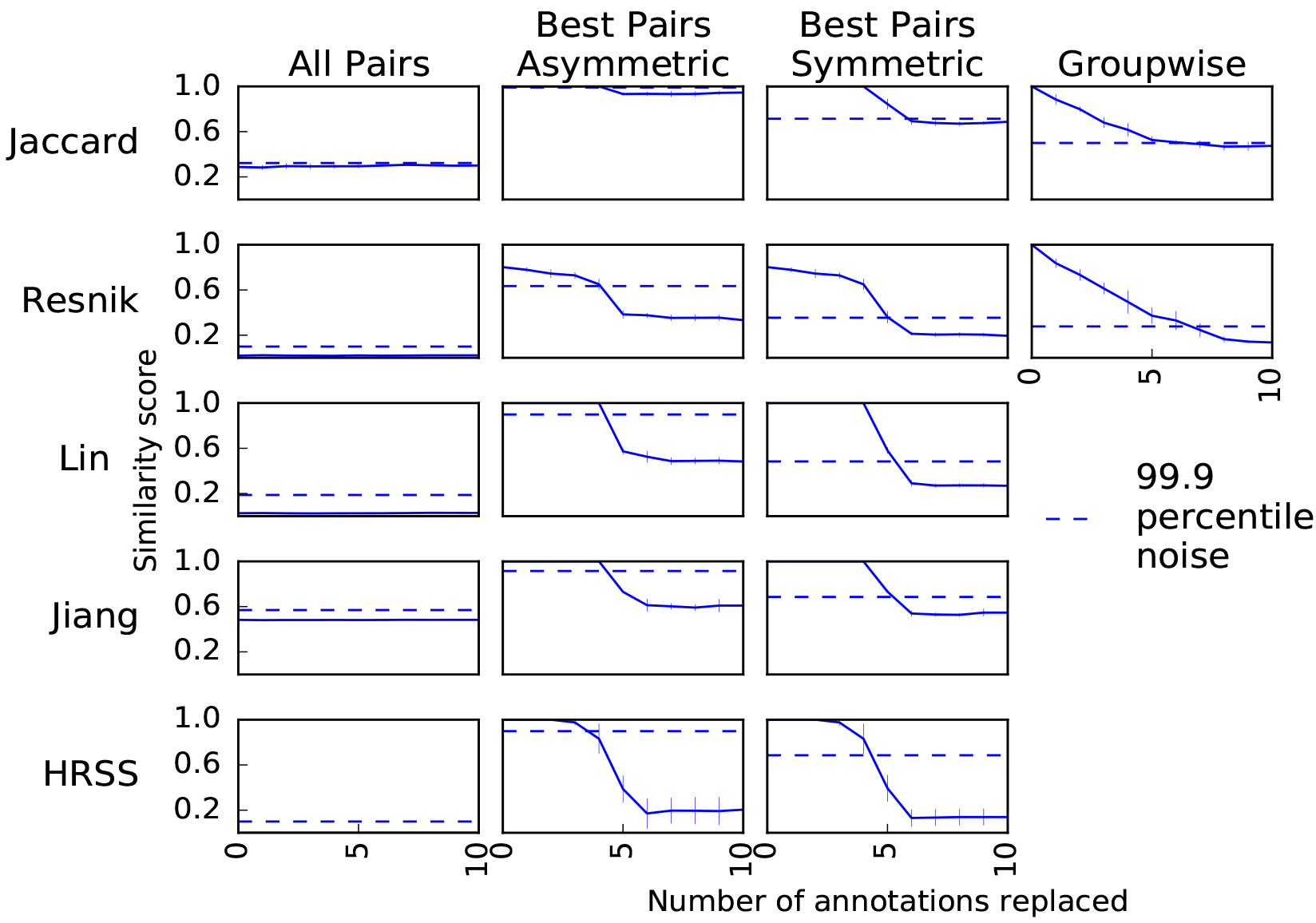
Pattern of similarity decay with five profiles of size 10 are decayed via Random Replacement. Similarity is computed using five semantic similarity metrics (Rows 1-5) in combination with four profile similarity methods (Cols 1-4). Solid lines represent the mean best match similarity of the five query profiles to the database after each annotation replacement. Error bars show two standard errors of the mean similarity across the five profiles. Dotted lines represent the 99.9th percentile of the noise distribution.

Groupwise measures (for Jaccard and Resnik) show a gradual decline in similarity unlike the sharp decline exhibited by Best Pairs methods. The magnitude of discrimination from noise is greater than for Best Pairs, and discrimination is still possible beyond 50% decay.

These results from the comparison of the four profile similarity approaches show that Best Pairs Symmetric (among pairwise statistics) and Groupwise result in the greatest sensitivity across the tested similarity metrics. Accordingly, these two profile similarity methods were selected for subsequent experiments.

Since Groupwise statistics are available only for two of the five similarity metrics, we focus only on Best Pairs Symmetric to compare the five semantic similarity metrics (Figure 4, Rows 1-5, Col 3). We observe that among the IC based measures (Resnik, Lin, and Jiang), Lin shows the greatest discrimination from noise. All three metrics decline into noise similarity at the 50% decay mark. Lin and Jiang show a flat performance until 50% decay before suddenly dipping below the noise threshold. This indicates that these metrics fail to distinguish between perfect similarity between identical profiles (no decay) from imperfect similarity between decayed profiles before 50% decay. On the contrary, Resnik displays a gradual decline in similarity indicating greater discrimination between true matches of varying quality. Comparing the distance based metric (Jaccard) to the hybrid metric (HRSS), HRSS declines below the noise threshold at the 30% decay mark unlike Jaccard which discriminates from noise until 50% decay.

Based on these results, we conclude that Resnik (among the IC-based metrics), and Jaccard (between distance-based and hybrid metrics) demonstrate the greatest sensitivity. These two metrics in conjunction with the two selected profile similarity methods were used for all further experiments.

### 4.3 Improving the sensitivity of Best Pairs metrics

The sharp decline in similarity under the Best Pairs statistics at approximately 50% decay can be understood as a result of summarizing pairwise similarity scores with the median of the distribution Manda et al. [2016]. To understand if the sensitivity of pairwise statistics such as Best Pairs can be tuned using a different percentile of the pairwise score distribution, we compared the results using the 80th percentile rather than the median.

Best Pairs Jaccard and Resnik distinguish similarity from noise for greater levels of decay when the 80th percentile is used (Figure 5) than for the median. This illustrates that sensitivity for pairwise metrics can be tuned by how the pairwise similarities are aggregated. Jaccard and Resnik perform similarly with respect to how long similarity can be distinguished from noisewith Groupwise showing marginally less sensitivity. We again see that Group-wise has a more gradual decline in similarity. The implication of this is that Groupwise statistics will provide more fine discrimination among matches of varying quality and thus be better for rank ordering the strength of matches Manda et al. [2016], while sensitivity may be slightly greater for Best Pairs when using a high percentile.

**Figure 5:**
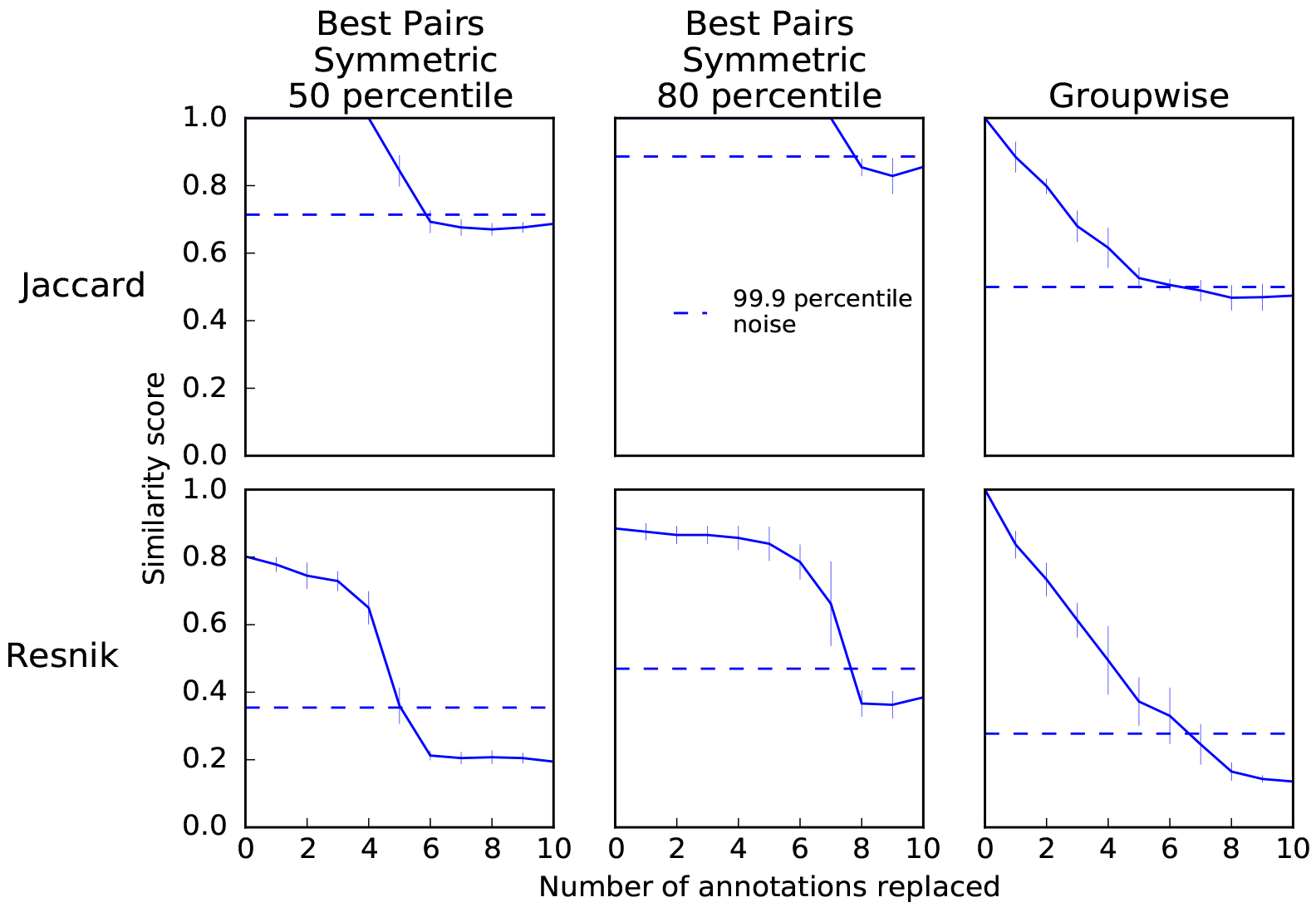
Comparison of the sensitivity of Best Pairs (summarized using the 50th percentile or the 80th percentile) and Groupwise metrics. Solid lines represent the mean best match similarity of the five query profiles to the database after each annotation replacement. Error bars show two standard errors of the mean similarity across the five profiles. Dotted lines represent the 99.9th percentile of the noise distribution.

**Figure 6:**
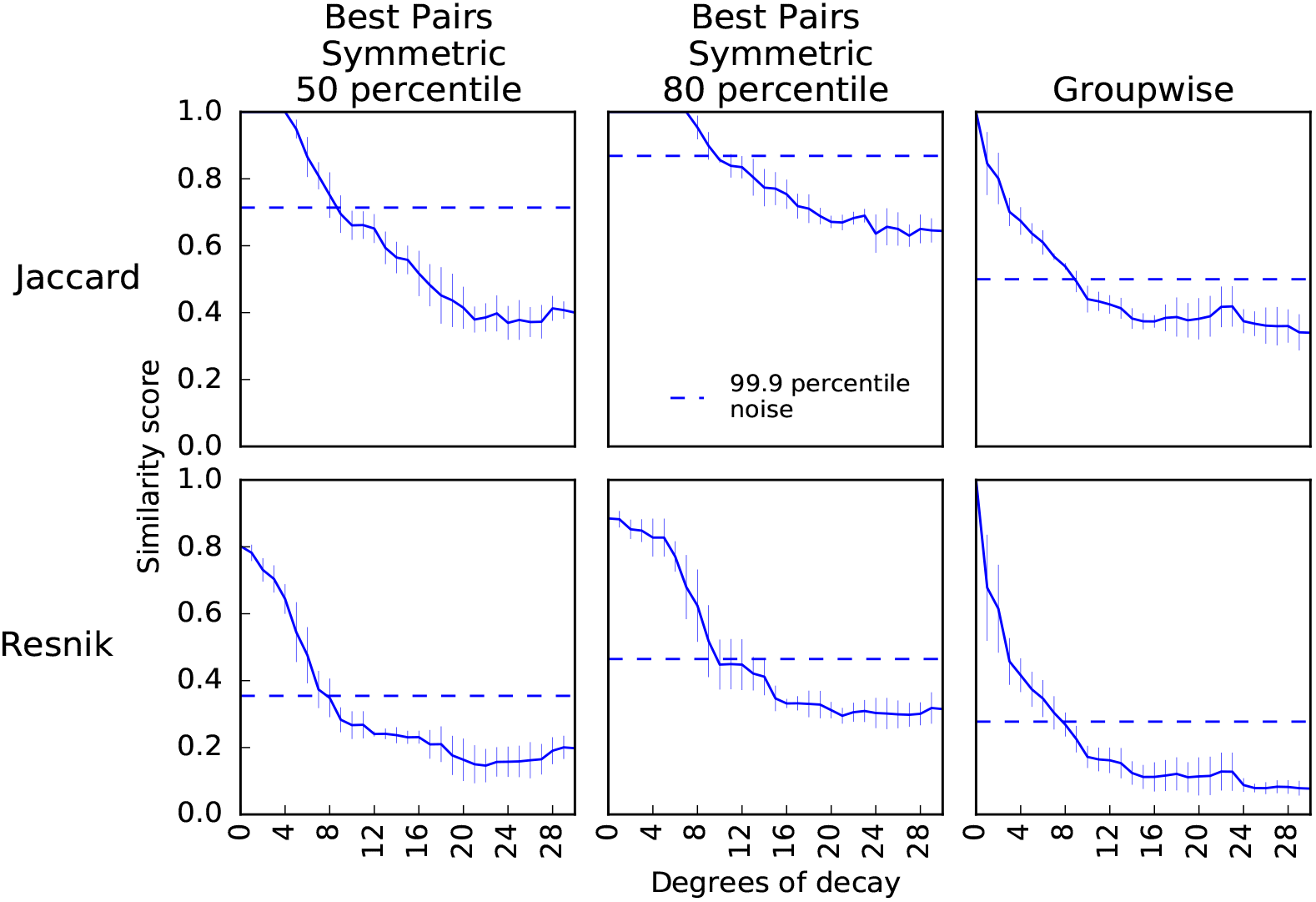
Pattern of similarity decay with the Ancestral Replacement model. Solid lines represent the mean best match similarity of the five query profiles to the database after each annotation replacement. Error bars show two standard errors of the mean similarity across the five profiles. Dotted lines represent the 99.9th percentile of the noise distribution.

### 4.4 Similarity Decay under the Ancestral Replacement Model

Next, we explored if the metrics exhibit the same relative performance when using the Ancestral Replacement decay mmodel rather than Random Replacement (Section 3.1.2).

The results for Ancestral Replacement are in general agreement with those reported above for the Random Replacement. Changing the percentile at which pairwise scores are aggregated again shows the percentage decay at which similarity for the Best Pairs statistics can no longer be discriminated from noise to be at the percentile used (either 50% or 80%). Groupwise metrics again, show a more gradual decline and fail to discriminate signal from noise at less extreme level of decay than for the Best Pairs statistic using the 80th percentile.

## 5 Discussion

Our findings reveal that sensitivity can vary dramatically among semantic similarity metrics and among different parameter choices. The majority of studies that use semantic similarity employ the Best Pairs or All Pairs approaches to aggregate similarity between two profiles, employing a variety of semantic similarity metrics. Here we see pronounced performance differences among these metrics. The way in which pairwise statistics are aggregated has easily predictable effects upon sensitivity. In some case, Groupwise approaches may be more sensitive, and generally show greater discrimination among levels of similarity above the sensitivity threshold. These results suggest specific ways to improve the sensitivity and interpretability of semantic similarity applications, particularly for profile comparisons.

We compared two models for decay of similarity between profiles, and found similar results for both. We also saw no substantive effect of profile size on the results. This increases our confidence in the generality of the results, although our evaluation is limited to one context, comparison among profiles sampled from among the Entity-Quality phenotype annotations in the Phenoscape KB.

Our results also illustrate how difficult it can be to statistically discriminate weakly matching profiles from noise, something which has received relatively little consideration in many applications of semantic similarity search to date. This suggests a need for more statistically informed reporting of results from semantic similarity matches, so that results which may be statistically meaningless are not interpreted as having biological significance.

## 6 Acknowledgements

We thank W. Dahdul, T.A. Dececchi, N. Ibrahim and L. Jackson for curation of the original dataset, along with the larger community of ontology contributors and data providers (http://phenoscape.org/wiki/Acknowledgments#Contributors), and useful feedback from P. Mabee, H. Lapp, W. Dahdul, and other members of the Phenoscape team. This work was funded by the National Science Foundation (DBI-1062542) and start-up funding to P. Manda from the University of North Carolina at Greensboro.

